# Simulations suggest robust microtubule attachment of kinesin and dynein in antagonistic pairs

**DOI:** 10.1101/2022.08.09.503394

**Authors:** Tzu-Chen Ma, Allison M. Gicking, Qingzhou Feng, William O. Hancock

## Abstract

Intracellular transport is propelled by kinesin and cytoplasmic dynein motors that carry membrane-bound vesicles and organelles bidirectionally along microtubule tracks. Much is known about these motors at the molecular scale, but many questions remain regarding how kinesin and dynein cooperate and compete during bidirectional cargo transport at the cellular level. The goal of the present study was to use a stochastic stepping model constructed by using published load-dependent properties of kinesin-1 and dynein-dynactin-BicD2 (DDB) to identify specific motor properties that determine the speed, directionality, and transport dynamics of a cargo carried by one kinesin and one dynein motor. Model performance was evaluated by comparing simulations to recently published experiments of kinesin-DDB pairs connected by complementary oligonucleotide linkers. Plotting the instantaneous velocity distributions from kinesin-DDB experiments revealed a single peak centered around zero velocity. In contrast, velocity distributions from simulations displayed a central peak around 100 nm/s, along with two side peaks corresponding to the unloaded kinesin and DDB velocities. We hypothesized that frequent motor detachment events and relatively slow motor reattachment rates resulted in periods in which only one motor is attached. To investigate this hypothesis, we varied specific model parameters and compared the resulting instantaneous velocity distributions, and we confirmed this systematic investigation using a machine learning approach that minimized the residual sum of squares between the experimental and simulation velocity distributions. The experimental data were best recapitulated by a model in which the kinesin and dynein stall forces are matched, motor detachment rates are independent of load, and the kinesin-1 reattachment rate is 50 s^-1^. These results provide new insights into motor dynamics during bidirectional transport and put forth hypotheses that can be tested by future experiments.

**Statement of significance:** Bidirectional transport of vesicles along microtubules is vital for cellular function, particularly in the highly elongated axons and dendrites of neurons, and transport defects are linked to neurodegenerative diseases. For developing future therapeutic strategies, a better understanding is needed for how motors, cargo adapters, and accessory proteins coordinate their activities to transport cargo to their proper cellular locations. We approached this problem by simulating how antagonistic kinesin and dynein motors compete in pairs. We constrain our simulations by recent experimental results and conclude that the motors spend nearly all their time attached to the microtubule and competing against one another. This behavior is not predicted by existing single-molecule experiments and thus provides new insights into bidirectional transport.

## Introduction

Kinesin and dynein motor proteins carry out anterograde and retrograde transport in cells (1,2) and work together to achieve long-distance bidirectional transport in neurons (1,3–6). Coordinated transport is important for neuron growth and function (7–9), and dysfunction can lead to neurodegenerative diseases such as ALS and Alzheimer’s (8,10,11). However, the mechanisms through which kinesin and dynein cooperate during cargo transport are unclear. Tracking of vesicles and other cargo in cells can reveal the complex dynamics of bidirectional transport (5,12–15), but this approach does not allow direct observation of the motors involved, and interpretations are complicated by the many regulatory factors that control intracellular transport (9). Single-particle tracking and optical tweezer studies have uncovered key details of the mechanisms by which individual motor proteins walk along microtubules (16–18), but apart from a few exceptions (19–23), experiments involving antagonistic motor pairs or teams are lacking. Computational simulations provide a valuable tool to bridge the gap between single-motor studies *in vitro* and cargo transport observations in cells, and these approaches help to uncover aspects of motor function that are difficult to observe through experiments.

A number of stochastic stepping models have been developed to simulate microtubule-based transport by kinesin and dynein motors (24–29). These models are parameterized based on single-particle tracking and optical tweezer experiments, and the simulation results can be used to investigate the influence of specific motor parameters on the resulting bidirectional transport directionality and speed. However, one challenge is that bidirectional cargo trajectories are inherently complex and involve both directional switching and fluctuating velocities, making it difficult to quantitatively compare simulations and experiments. A second challenge is that simulation results are highly dependent on choices of specific parameters that describe motor stepping and motor-microtubule binding/unbinding kinetics, and in many cases these parameters are not tightly constrained by existing experiments.

There have been several experimental developments over the last few years that motivate the next generation of bidirectional stepping models. The first is the appreciation that traditional single-bead optical tweezers create non-negligible vertical forces normal to the microtubule that can accelerate the detachment of kinesin under load (30). This effect was clearly demonstrated by the finding that, compared to the single-bead assay, the kinesin-1 attachment duration at stall increases substantially in a three-bead assay, where vertical forces are eliminated (31). The second major development is the finding that cytoplasmic dynein activated by cargo adaptors such as BicD2, BicDR1, and Hook3 is highly processive and can generate forces in the range of kinesins (32–39). A third important discovery was that activated dynein complexes such as dynein-dynactin-BicD2 (DDB) frequently switch between three motility states – processive, paused, and diffusive – when engaged with a microtubule (34,37,40). Computational simulations provide an important tool to unravel how these different factors play into the bidirectional transport achieved by pairs or teams of kinesin and dynein motors.

The goal of present work is to incorporate recent kinesin and DDB experimental insights into a stochastic stepping model that recapitulates the bidirectional transport behavior of single kinesin-1 and DDB motor pairs. Parameter sensitivity tests and an objective machine learning approach explored the influence of motor detachment/reattachment dynamics and stall force parameters on the resulting bidirectional transport dynamics. The simulations were tuned to match recent in vitro experiments that tracked the dynamics of kinesin-DDB (Kin-DDB) pairs connected through complementary DNA hybridization. We found that incorporating detachment and reattachment rates inferred from published work resulted in directional switching and fast plus- and minus-end velocities not observed in the experiments. Instead, experimental data were best recapitulated by a model that incorporated matched stall forces and load insensitive detachment for both kinesin-1 and DDB. Thus, these simulations predict that motors working in antagonistic pairs have different properties than motors in isolation, and these properties enhance competition between kinesin and dynein.

## Methods

### Stochastic stepping algorithm

Kin-DDB bidirectional transport was simulated using an updated and modified version of a previously published model (27) consisting of one kinesin and one dynein attached to a virtual cargo. At each timepoint, any attached motor can step forward by 8 nm, step backward by 8 nm, or detach from the microtubule; any detached motor can reattach to the microtubule. In the DDB switching model, an attached DDB motor can also switch between processive, diffusive, and stuck states. The decision for what event occurs and the time of the transition is decided by time evolution of the system using the Gillespie Stochastic Simulation Algorithm (41), as follows. For an event with a first-order transition rate constant, *k*, the transition time is generated as:

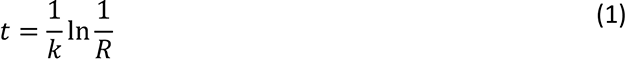

where R is a uniformly distributed random number in range 0 to 1. In a system with N possible events, the rate of any event occurring equals the sum of the rate constants for all possible events. Thus, at each time point, a two-stage process (the ‘direct method’ of Gillespie (41)) was used to determine the time and identity of the next transition. First, a random number *R*_1_ is generated and used to compute the time to the next event, *i* as follows:

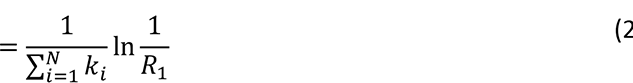

In the second stage, a new round of random number generation is used to determine which of all possible events occurs; here, the probability of any event occurring is proportional to its rate constant (27,41). We split this stage into two steps, with the goal of modularizing the code and enabling expansion to multiple motors bound to the same cargo. We define *j* = 1,…M motors in the simulation (for our Kin-DDB simulations, M = 2). For each motor, *j*, there are *i* = 1,…N possible transitions, where N = 4 in most cases for our model (forward step, backward step, attachment and detachment). In the first step, a uniformly distributed random number, *R*_2_ is generated and used to choose which of the M motors present will make a transition, and in the second step a random number *R*_&_ is used to choose which of the N possible transitions for that motor will occur. Formally, using M motors, the probability of motor *j*, which has N possible transitions acting is:

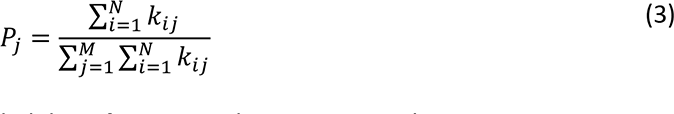

From N possible events on motor j, the probability of event i with rate constant kjj occurring is:

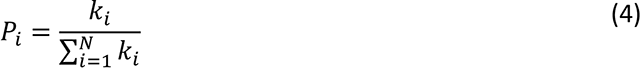

Where 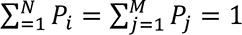. To identify the next event, the unit interval is divided into segments corresponding to each possible transition: 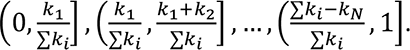. A random number is then used to determine which transition will occur. This method is repeated until all motors detach from the microtubule or a maximum run length or run duration is reached.

In the simulations, force is defined as positive in the plus-end direction and negative in the minus-end direction; thus, for kinesin-1 negative forces are hindering loads and positive forces are assisting loads. The force, *F*, applied to each motor is defined as the motor-cargo distance, Δ*x*, multiplied by stiffness of the motor, *k*_*motor*_:

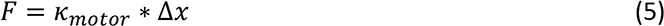

In our previous model (27), we used *k*_*sin*_ = 0.3 pN/nm and *k*_*DDB*_ = 0.065 pN/nm based on published work (42–44). For investigating the role of stiffness in the present work, we simplified the model by incorporating an identical stiffness, *k*_*motor*_ = 0.1068 pN/nm for each motor, which sets the cargo at the midpoint between two motors.

At every timepoint, the position of the virtual cargo is set by computing a force balance between the two motors when both motors are attached, and by setting it to the position of the attached motor if only one motor is attached. Thus, following a motor step or detachment, the time for the system to reach mechanical equilibrium is assumed to be negligible. This instantaneous force balance can be justified by considering the relaxation time of the ∼30 nm quantum dot attached to the Kin-DDB complex in the experiments. The drag coefficient γQdot of a microsphere with radius rQdot = 15 nm in an aqueous solution (viscosity coefficient η = 10^-9^ pN·s/nm^2^) is given by the Stokes’ law as γQdot = 6πηrQdot = 2.8 × 10^-7^ pN·s/nm (45). Following a motor step, we can calculate the characteristic relaxation time constant for the particle to exponentially relax back to an equilibrium position as *τ* = *γ*_*Qdot*_/*k*_*motor*_, where *k*_*motor*_ = 0.1068 pN/nm. This calculation yields a 3-microsecond relaxation time constant, which is negligible compared to the motor stepping and attachment/detachment rates in the msec range or slower.

### Kinesin-1 simulation parameters

The kinesin-1 stepping model incorporates a linear force-velocity relationship with a constant backstepping rate, based on published optical tweezer measurements (46,47). The kinesin unloaded velocity, *v*^6^ = 515 nm/s (Fig. S1A) was determined by carrying out single-molecule total internal reflection fluorescence experiments in the buffer used for the published Kin-DDB experiments (30 mM HEPES, 1 mM EGTA, 50 mM K-acetate, 2 mM Mg-acetate, 10% glycerol, 1 mM ATP, 10 μM Taxol, 0.2 mg/ml casein, pH 7.4) (34). The motor velocity is equal to the net stepping rate multiplied the 8 nm step size, *v* = *k*_forward_ − *k*_back_: ∗ 8 *nm*. Under assisting loads, the motor velocity is constant and equal to the unloaded velocity (46,47). Under hindering loads the velocity decreases linearly to zero at the stall force (*F*_*stall*_ = 8 pN), where the forward and backwards stepping rates are equal, and plateaus at the constant and load-independent backstepping rate, *k*_*back*_ of 3 *s*^,1^ (46,47). Based on the measured unloaded velocity and the constant backstepping rate, the unloaded forward stepping rate, *k*^0^_*forward*_ is defined as:

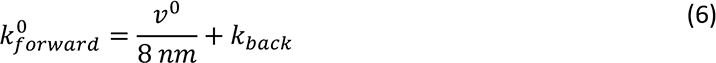

Under hindering loads up to the limit where *k*_*forward*_(*F*) = 0, the load-dependent forward stepping rate is:

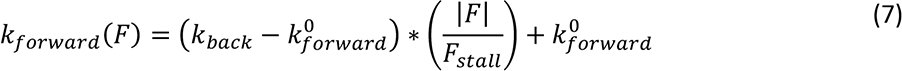

### The kinesin-1 microtubule detachment rate *k*_*detach*_(*F*) varied exponentially with load (F)

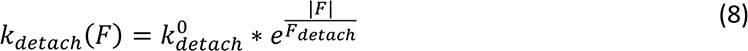

Here, is the unloaded detachment rate, and the detachment force parameter 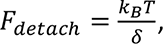 where *k*_*B*_is Boltzmann constant, T is absolute temperature, and *δ* is a distance parameter that defines the load-sensitivity of detachment. Based on Andreasson et al. (47), the unloaded detachment rate, *k*^6^ and the detachment force parameter, *F*_*detach*_ were set to 1.11 *s*^,1^ and 6.83 pN under hindering loads, and 7.4 *s*^,1^ and 12.8 pN under assisting loads, respectively.

### DDB simulation parameters

The DDB kinetic model is based on load-dependent velocity measurements by Elshenawy et al. (33), with simplifications to enable more straightforward parameter sensitivity tests. The DDB step size in both directions was set to 8 nm, the stall force was set to 3.6 pN, and a constant and load-independent backstepping rate, *k*_*back*_ = 15 *s*^,1^ was included (33). The unloaded velocity of DDB was set to 328 nm/s based on published control experiments carried out in parallel with the Kin-DDB experiments (34). Under hindering loads (which are positive for DDB), a linear DDB force-velocity curve was used, similar to Eq. 7 for kinesin, up to the force at which *k*_*forward*_(*F*) reached zero. Under assisting loads (which are negative for DDB and occur only in rare instances where DDB is positioned to the plus-end of kinesin-1), the load-dependent DDB velocity was taken from published work (33):

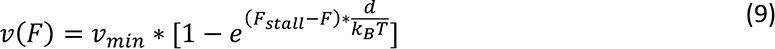

Here *v*_*min*_ is the asymptotic velocity under super stall force, equal to -201 nm/s, and *d* is the characteristic distance of 1.5 nm that defines the load-dependence of the stepping rate (33). In these assisting load cases, the load-dependent forward stepping rate, *k*_*forward*_(*F*) was calculated as:

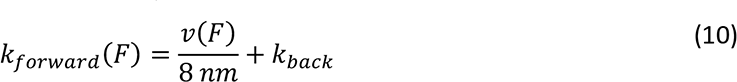

The DDB detachment rate, *k*_*detach*_, was modeled with an exponential load dependence similar to kinesin (Eq. 8), with *k*^6^ of 0.1 s^-1^ and *F*_*detach*_ of 3 pN, based on experimental work from Belyy et al. (32).

### DDB state-switching model

In a subset of simulations, we incorporated DDB state switching based on work from Feng et al. showing that DDB switches between processive, diffusive and stuck states during movement (34). To simulate this behavior, three motility states for DDB were integrated into the model, as follows. In the processive state, DDB can move forward, backward, or detach; in the stuck state, DDB cannot move or detach from the microtubule; and in the diffusive state, DDB offers no resistance to kinesin movement and can also detach from the microtubule. From every state, the motor can transition to either of the remaining states, with transition rates as follows: the processive-to-stuck switching rate was 1.0 *s*^,1^ with reverse rate of 1.8 *s*^,1^; the stuck-to-diffusive switching rate was 0.07 *s*^,1^ with a reverse rate of 0.33 *s*^,1^; and the diffusive-to- processive switching rate was 3.9 *s*^,1^ with a reverse rate of 0.23 *s*^,1^ (34). In the state-switching model simulations, these transitions were added to the stepping and attachment/detachment transition possibilities for DDB. For the majority of the simulations, this state switching was turned off.

### Data processing

After running the simulations, the raw data were processed to match the experimental conditions (34), as follows. To match the 20 fps frame rate (34), cargo position was averaged over 50 ms windows. To account for uncertainties in fitting the experimental point spread function, a normally distributed error with an 8 nm standard deviation was added to cargo position at each time point. The instantaneous velocity at each timepoint was calculated by taking three-point slope of positions 50 ms before and after each point.

### Parameter optimization

As an alternate approach for parameter identification, we used a Bayesian Optimization method for minimizing the residual sum of squares (RSS) between the experimental and model instantaneous velocity distributions. The instantaneous velocity probability density functions from -2000 nm/s to 2000 nm/s with 10 nm/s bin widths were calculated for both experimental data and simulation results, where *P*_i_ is the PDF of instantaneous velocity *i*. For each parameter set, a velocity probability density function was generated from 1000 trajectories having a maximal run time of 50 s. The Matlab function ‘bayesopt’ was used with 100 iterations to identify parameter sets that minimized the RSS function:

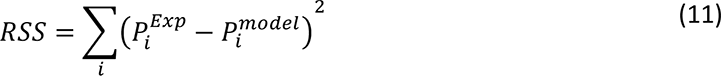

## Results

### Formulation of bidirectional stepping model

In recently published experiments, we reconstituted bidirectional cargo transport *in vitro* by connecting a truncated kinesin motor to an activated dynein-dynactin-BicD2 (DDB) complex through complementary single stranded DNA oligonucleotides (34). A quantum dot was then linked to a biotin on one end of the DNA (Fig. 1A), and the position of the fluorescent cargo was tracked via total internal reflection fluorescence (TIRF) microscopy at 20 frames per second. Consistent with previous work (33), the resulting complexes moved slowly for long durations along immobilized microtubules, with some complexes moving net plus-end and others moving net minus-end (Fig. 1C) (34). To better understand the kinesin and dynein motor dynamics underlying this bidirectional transport, we adapted a previously published Kin-DDB transport bidirectional transport model, carried out simulations, and compared predictions from the simulations to the new experimental data. The model uses the Gillespie Stochastic Simulation Algorithm (41), as described in Methods and our previous publication (27). At every time point, either motor can step forward, step backward, detach from or reattach to the microtubule, with the probability of each transition being proportional to its first-order rate constant (Fig. 1B; diagram of the algorithm is given in Fig. S2). The simulated cargo has neither mass or drag, and its position is updated following every step to maintain force balance. The parameters for this ‘basic model’ are shown in Table 1.

**Figure 1.**
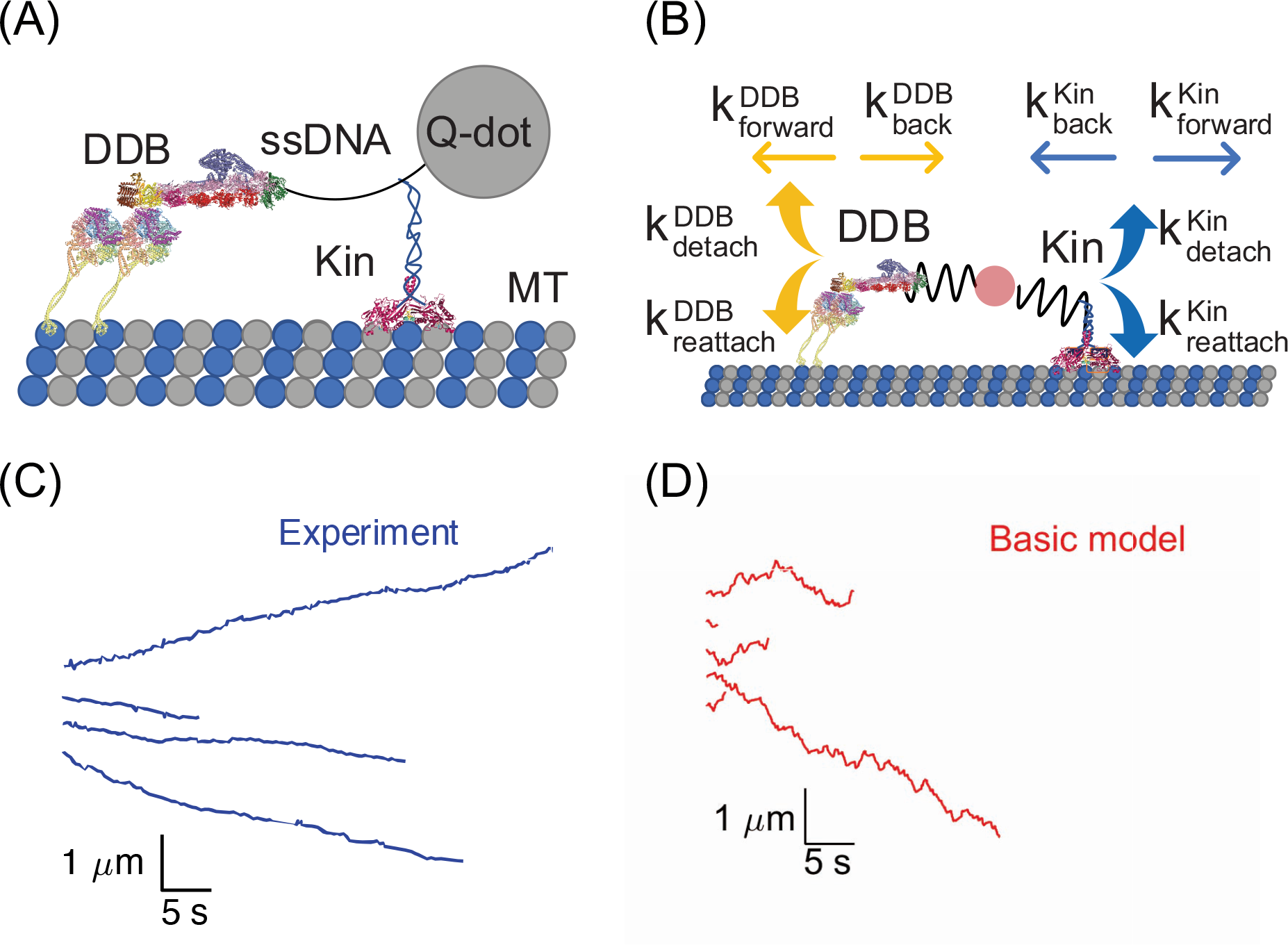
(A) Schematic of Kin-DDB bidirectional transport tracking experiment in which motors are connected together by a complementary single stranded DNA (ssDNA) and attached to a quantum dot for visualization by fluorescence microscopy (34). (B) Stochastic model of Kin-DDB bidirectional transport showing the different rate constants incorporated into the simulations. (C) Example individual distance versus time traces of experimental data from published work by Feng et al (34). (D) Simulation traces from the basic model. Kinesin and DDB images adapted from PDB files 6A1Z and 3VKH (49–51).

**Table 1.**
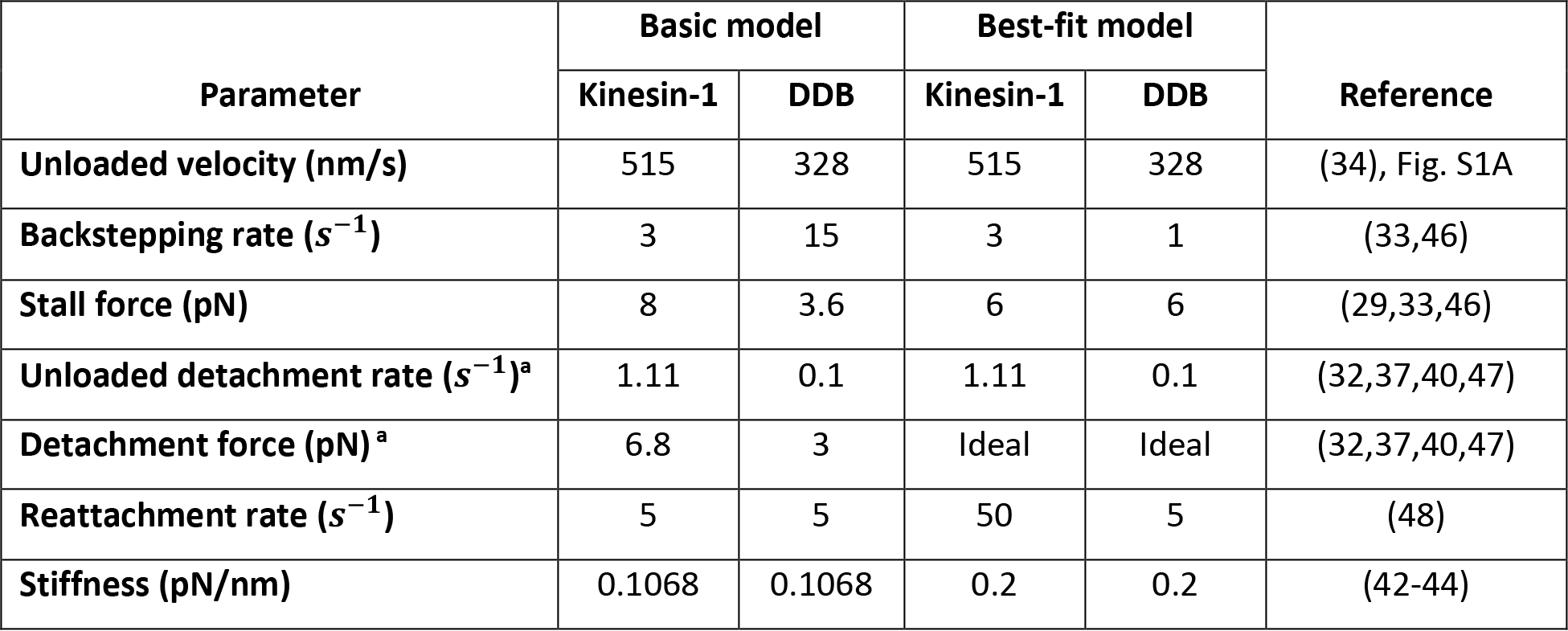
Parameters used for the initial ‘basic’ model and the final ‘best-fit’ model. Parameters for the ‘better-fit’ model are given in Table S1. ^a^ *For kinesin under assisting loads, the unloaded detachment rate extrapolation was 7.4 s^-1^ and detachment force was 12.8 pN based on* (*47*)*. For DDB under assisting loads, the unloaded detachment rate and detachment force were identical to the hindering load condition. Ideal bond (load-independent detachment) corresponds to Fdetach equal to infinity*.

### Comparing bidirectional stepping simulations to experimental data

We next simulated bidirectional stepping of a complex containing one kinesin-1 and one DDB motor to determine the degree to which the simulations can recapitulate the experimental results (Fig. 1C). The mean velocities of trajectories from the simulations (-4.4 ± 115 nm/s mean and standard deviation; N = 1000) were similar to the experimental velocities (-8.2 ± 47 nm/s mean and standard deviation; N = 30). However, the simulated trajectories contain fluctuations and directional switching not seen in the experiments (Fig. 1C and D). One explanation for the larger fluctuations in the simulated traces is that experimental temporal and spatial resolution limits are obscuring fluctuations in the experimental data. Therefore, to enable more accurate comparison between experiments and simulations, we added simulated experimental noise and reduced the temporal resolution to the simulations, as follows. From the quantum dot tracking experiments (34), we determined the experimental error in the Gaussian fits to the point-spread function data to be 8 nm; thus we added a normally distributed noise term with a standard deviation of 8-nm to each point in the simulation. Second, we binned the simulated data to 50 ms to match the 20 per second frame rate of the experiments.

To better compare the fluctuations in the traces resulting from motor stepping dynamics, we examined 5-second segments of experimental and simulated traces. Despite the downsampling, larger and more frequent fluctuations can be seen by eye in the simulated traces compared to the experimental traces (Fig. 2A and B). A Mean Square Displacement (MSD) analysis quantitatively confirmed this: simulated traces had an apparent diffusion coefficient of 11,800 *nm*^2^/*s*, compared to 996 *nm*^2^/*s* for the experiments (Fig. S1B). To better compare the experimental and simulated velocities, we calculated the instantaneous velocities over 100-ms time windows for both the experimental and simulation results and plotted the instantaneous velocity distributions (Fig. 2C). The experimental instantaneous velocity distribution had a prominent single peak centered at -12 nm/s, and 95% of the distribution was between -394 and 371 nm/s. In contrast, the simulated instantaneous velocity distribution had a distinct central peak that was right shifted compared to the experiments, along with distinct side peaks corresponding to the unloaded velocities of DDB and kinesin-1. Using a Gaussian mixture model (GMM), the velocity distribution was fit well by three normal distributions: a peak centered at 127 nm/s that accounted for 53% of the population, a minor peak at -334 nm/s that accounted for 41% of the population, and another minor peak centered at 587 nm/s that accounted for 6% of the population (Table 2). The motor reattachment rate in our simulations was 5 s^-1^ based on published experimental data for kinesin-1 (48,52,53), and because equivalent experimental data are not available for dynein, we chose the same value for the DDB complex

**Figure 2.**
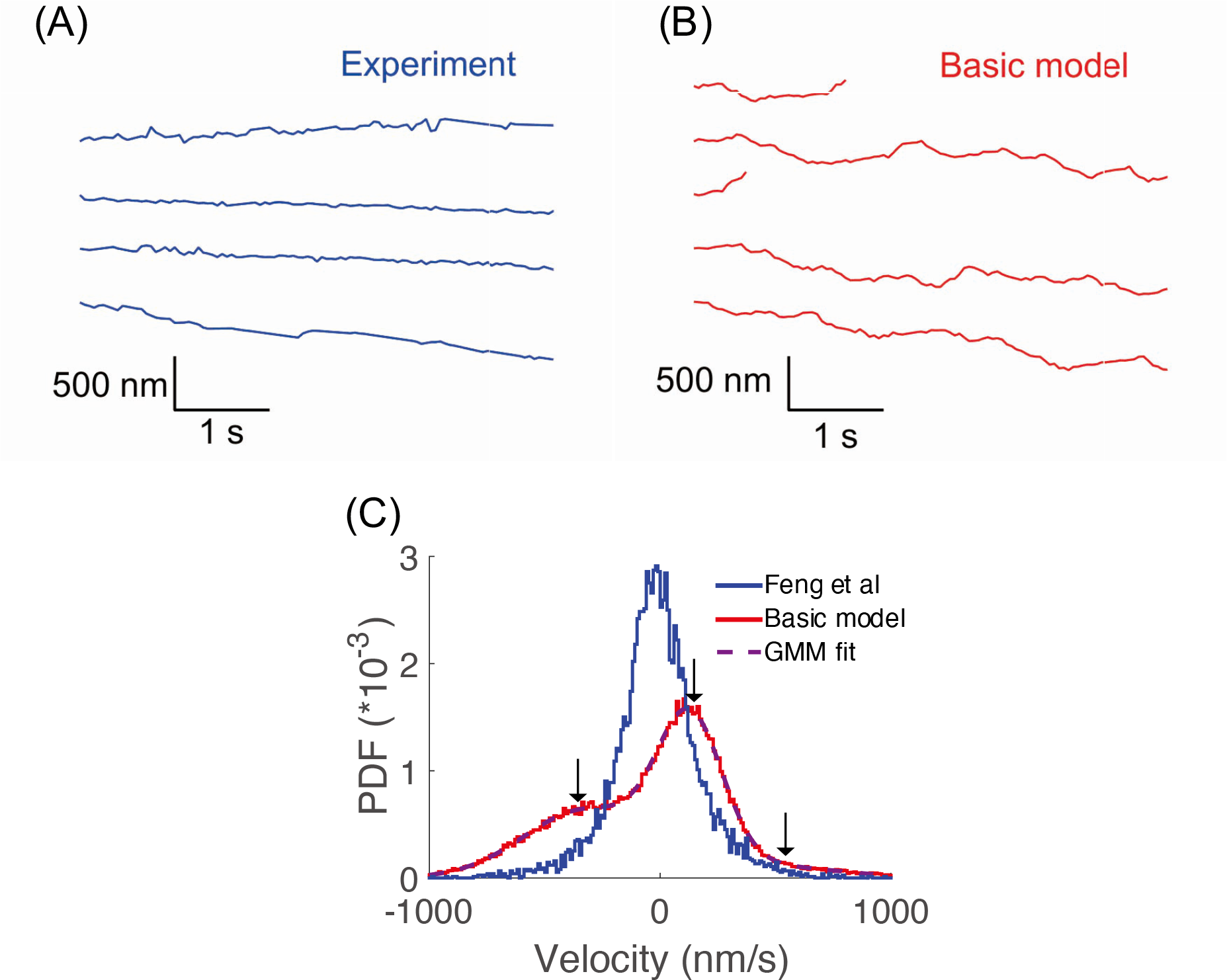
Comparison between simulation results and experimental data from Feng et al. (34). (A and B) Example individual distance versus time traces over a 5-second window, showing that simulation traces contain more fluctuations than the experimental traces. Model traces were downsampled to 20 Hz and positional noise was added to more accurately match experimental conditions. (C) Instantaneous velocity distribution (averaged over 100 ms windows with 10 nm/s bin width), showing three peaks in the simulation results (red, N = 1000 traces), but only a single central peak in the experimental distribution (blue, N = 30 traces). Parameters for Gaussian mixture model (GMM) fit are given in Table 1.

**Table 2.** *Gaussian Mixture Model fits to instantaneous velocity distributions. Experimental data from Feng et al.* (*34*) *include N = 30 traces and basic and best-fit models both include N = 1000 traces. Percentages are the relative integrated weights of each mode. Fits used the “fitgmdist” function in MATLAB with three Gaussians having initial values of 0, 500, and -300 nm/s). RSS values denote the residual sum of squares between the probability density functions each model and the experimental data*.

|  | Experiments | Basic model | Best-fit model |
| --- | --- | --- | --- |
| **Central peak** | -18 nm/s (65%) | 127 nm/s (53%) | -8 nm/s (93%) |
| **Plus-end peak** | 98 nm/s (23%) | 587 nm/s (6%) | 452 nm/s (0.5%) |
| **Minus-end peak** | -204 nm/s (11%) | -334 nm/s (41%) | -319 nm/s (6.5%) |
| **Residual sum of squares (RSS)** | | $3.93 \cdot 10^{-5}$ | $2.76 \cdot 10^{-6}$ |

(Table 1). This relatively slow reattachment rate means that when either motor detaches from the microtubule, it will take on average 200 ms to reattach, giving the other motor time to move at its unloaded velocity. Thus, these lateral peaks in the simulated velocity distribution can be explained by the simulations having more periods when only one motor is attached than the experiments, and their positions are close to the unloaded kinesin and DDB velocities.

To better understand the underlying motor dynamics that lead to the experimental Kin-DDB traces being relatively smooth and the velocity distribution having a single peak centered around zero, we investigated how the modeled properties of kinesin-1 and DDB contribute to the distinctive instantaneous velocity distribution of our ‘basic’ model described above. We hypothesize that there are properties of kinesin-1 and/or DDB motors that differ when the motors are joined in a two-motor complex compared to in their isolated states in the single-molecule fluorescence and optical trapping experiments used to develop the basic model. To test this overarching hypothesis, we varied motor parameters and quantified the change in performance, with the goal of reducing the magnitude of the fast plus- and minus-end velocity peaks in the instantaneous velocity histogram (Fig. 2C) to better match the experimental results.

### Testing the influence of dynein motor properties on bidirectional transport

The first hypothesis we tested was that the simulations differ from the experiments because the motile properties of DDB in a bidirectionally moving Kin-DDB complex differ from the DDB properties measured in single-motor optical tweezer experiments. To test this possibility, we first investigated whether either including experimentally observed pausing and diffusive states of the DDB complex could better align the simulations with the experiments. In virtually all published unloaded single-molecule studies, DDB exhibits long processive runs, but also undergoes episodes of 1D diffusion along the microtubule and spends a significant fraction of the time stuck to the microtubule in an immobilized state (32,34,37,38). Feng et al found that DDB spent 65% of the time in a processive state, 31% of the time in a stuck state, and 4% of the time in a diffusive state, and also quantified the switching rates between states (34). We incorporated this DDB switching behavior as a series of transitions into our model. We assumed that the diffusive state of DDB offered no resistance to kinesin stepping, and that in the stuck state, DDB neither moved nor detached from microtubule. In the instantaneous velocity distribution, incorporating this DDB switching behavior caused a decrease in the magnitude of the unloaded DDB velocity peak, but had no effect on the unloaded kinesin velocity peak (Fig. 3A). The fall in the unloaded DDB peak can be explained by DDB spending time in the stuck state instead of processive walking. The lack of change in the unloaded kinesin peak is likely due to the fact that DDB is in the diffusive state only a small fraction of the time (Fig. 3A). Thus, DDB state switching cannot account for the discrepancy between model and experimental results. Due to this lack of an effect, all subsequent simulations used only a single processive state for DDB.

**Figure 3.**
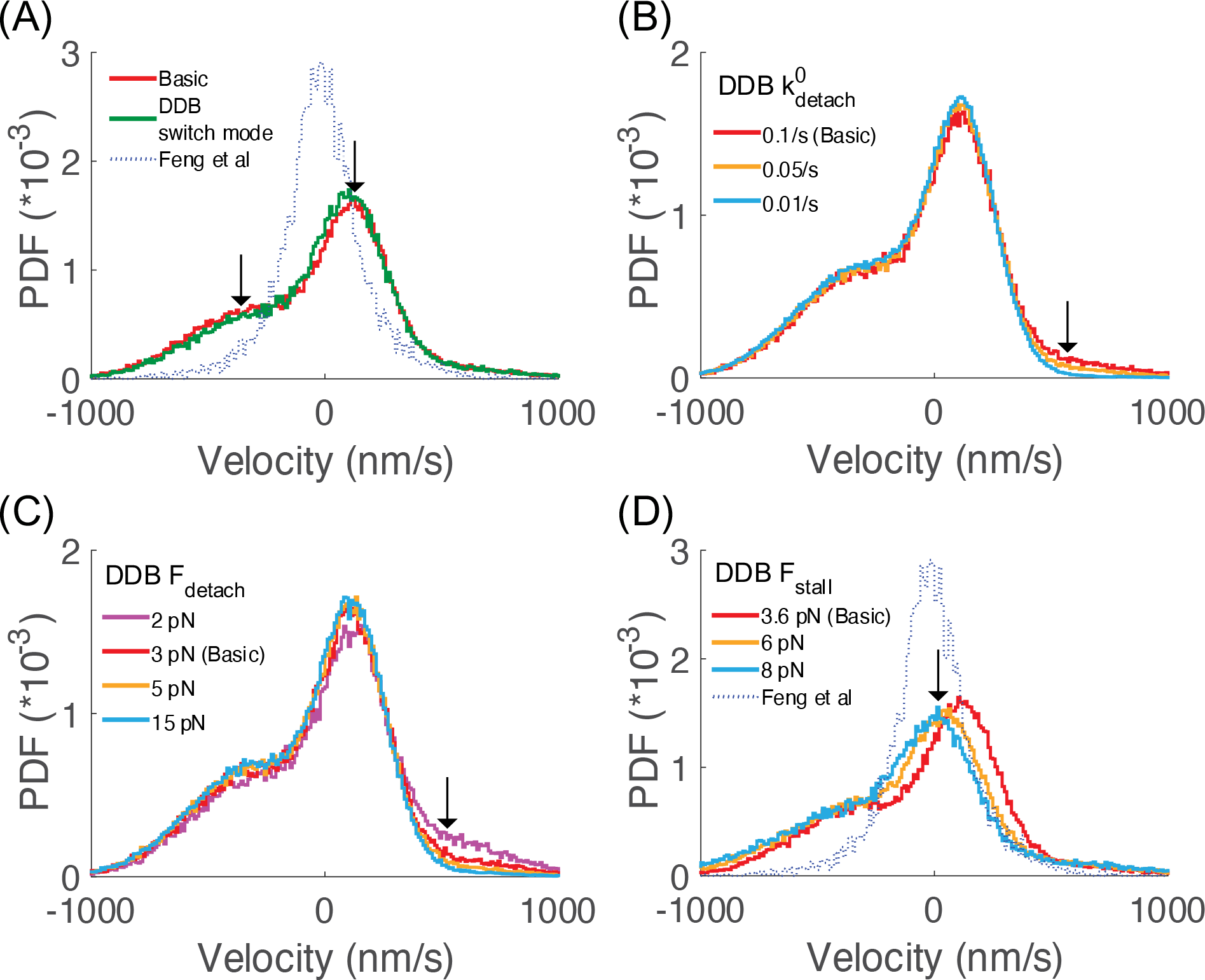
Effect of changing DDB parameters on the instantaneous velocity distribution. (A) Effect of incorporating switching between processive, diffusive and stuck states for DDB. Arrows denote the small changes in the central velocity peak and the minus-end side peak. (B) Effect of changing the DDB unloaded detachment rate, with arrow highlighting the resulting changes in the plus-end side peak. (C) Effect of changing the DDB detachment force, where larger Fdetach denotes less load-sensitivity of detachment. (D) Effect of increasing the DDB stall force, with arrow denoting leftward shift of the central peak with increasing stall force.

The second property we examined was the load-dependent detachment dynamics of DDB. There is abundant evidence in the literature that dynein alone has catch-bond behavior, meaning that the detachment rate slows down with increasing load (26,29,54–57). There is no experimental evidence to our knowledge that activated dynein in a DDB complex shows catch-bond behavior; however, only one published study directly addresses this question (33). Thus, we investigated whether the simulated bidirectional behavior is shaped by the load-dependent detachment kinetics of DDB by using an exponential dependence of the off-rate on load (Methods, Equation 8). In this formulation, a positive detachment force parameter (*F*_*detach*_) corresponds to a slip-bond, a negative value corresponds to a catch-bond, and a value of infinity corresponds to an ideal-bond. We found that slowing the unloaded detachment rate (*k*^6^) or increasing the detachment force (*F*_*detach*_) decreased the peak corresponding to the unloaded kinesin-1 velocity, bringing the distribution more in line with the experimental (Fig. 3B and C). When an ideal bond was simulated by setting *F*_*detach*_ to infinity, the instantaneous velocity distribution was indistinguishable to that for *F*_*detach*_ = 15 pN (Fig. S3A). Setting *F*_*detach*_ to -3 pN, corresponding to a catch-bond resulted in no change from the ideal-bond model (Fig. S3A).

The third property we examined was the DDB stall force, Fstall. In the literature, stall forces for isolated dynein were measured to be 1 pN by a number of labs (29), but activated dynein in a DDB complex was later shown to have a stall force of 3.6 pN (33). In our basic model, the kinesin stall force is twice that of DDB (Table 1) and the central peak in the instantaneous velocity distribution is shifted towards the plus-end by nearly 150 nm/s relative to the experimental peak (Table 2; Fig. 2C); thus, it is reasonable to expect that bringing the stall forces more into alignment should shift the central peak. We found this to be the case (Fig. 3D) – increasing the DDB stall force to 6 pN shifted the central peak leftward and increasing it to 8 pN to match the kinesin stall force brought the central peak nearly into alignment with the experimental peak. As a final investigation, we examined the sensitivity of the simulations to the DDB reattachment rate by increasing *k*_*attach*_ from 5 s^-1^ to 50 s^-1^. This modification had virtually no effect on the simulated velocity distribution (Fig. S3B). This lack of an effect likely stems from the slow DDB unloaded detachment rate of 0.1 s^-1^ – because DDB detachments are relatively infrequent, the unbound episodes are rare and shortening their duration has little effect.

To summarize the DDB parameter investigations, removing the load-dependence of DDB detachment by modeling DDB as an ideal-bond nearly eliminated the kinesin peak in the instantaneous velocity distribution, and increasing the DDB stall force shifted the central velocity peak leftward, closer to the experimental peak. In contrast, incorporating a switching model for DDB or increasing the DDB reattachment rate had only minimal effects on the velocity distribution. Next, to explore other modifications of the model that could diminish the DDB velocity peak, we examined the sensitivity of the model simulations to changes in the kinesin detachment and reattachment rates.

### Testing the influence of kinesin-1 motor properties on bidirectional transport

The second hypothesis we tested was that the discrepancy between simulations and experiments is due to differences in the motor properties of kinesin when operating in an antagonistic motor pair compared to single-bead optical tweezer experiments with isolated kinesin motors. These differences could be due to, for instance, the different geometries of being attached to a trapped bead versus being tightly connected to dynein through a short DNA (31,58,59), and they could in principle affect both motor on- and off-rates. The first investigation was to determine whether decreasing the detachment rate of kinesin by reducing *k*^6^ or increasing *F*_*detach*_ reduced the magnitude of the DDB velocity peak in the instantaneous velocity distribution. Decreasing *k*^6^ for kinesin-1 substantially decreased the magnitude of the unloaded DDB velocity peak and increased the magnitude of the central peak, although it also increased the unloaded kinesin velocity peak (Fig. 4A). Increasing *F*_*detach*_ to reduce the sensitivity of detachment to load had a similar effect but to a lesser degree (Fig. 4B), with the effect plateauing around an *F*_*detach*_value of 35 pN. Thus, decreasing the kinesin detachment rate diminished the minus-end velocity peak, bringing the simulations closer to the experimental results, but neither of these changes had a significant effect on the unloaded kinesin peak.

**Figure 4.**
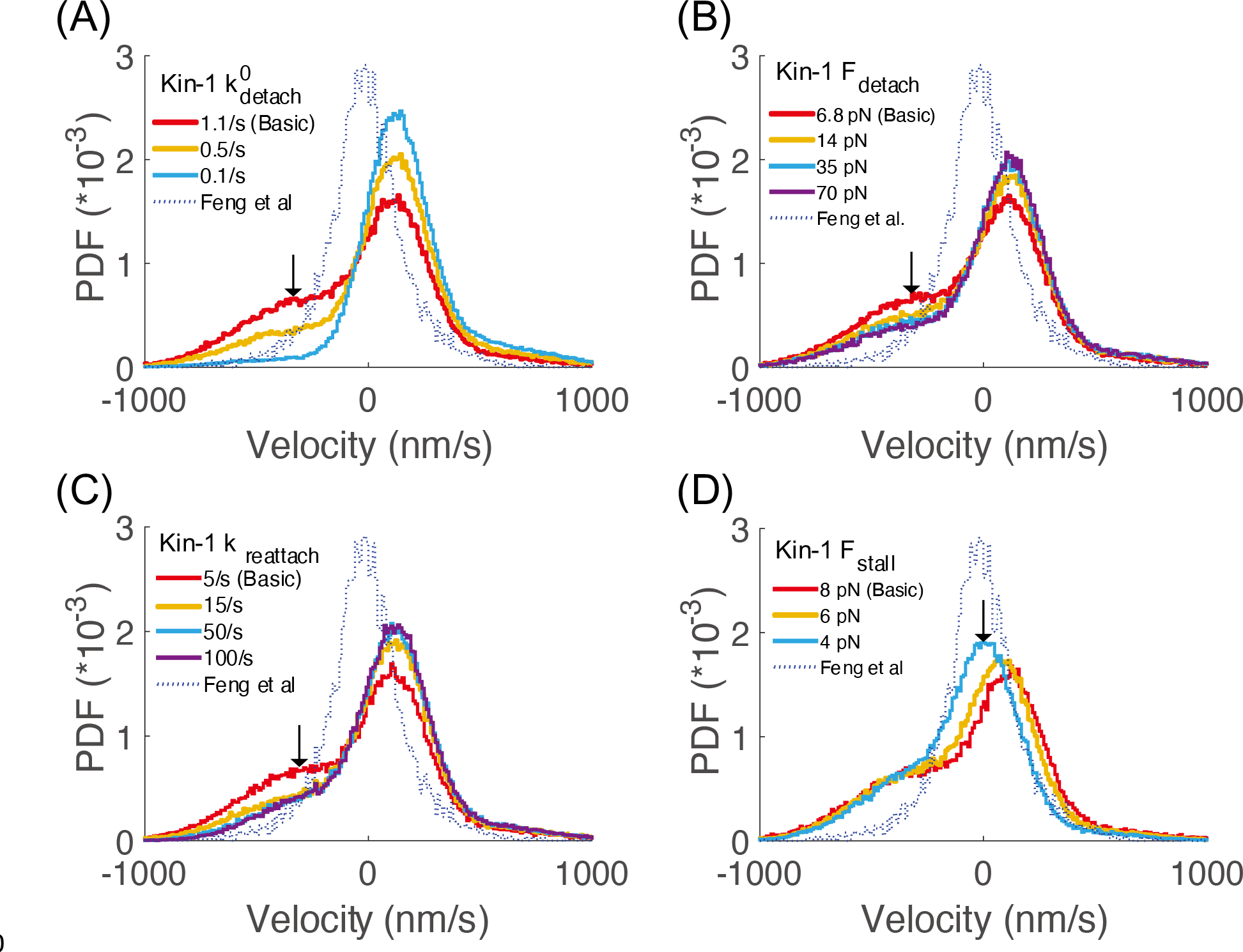
Effect of changing kinesin-1 parameters on the instantaneous velocity distribution. (A) Effect of decreasing the unloaded kinesin-1 detachment rate was to diminish the minus-end velocity peak and increase the central peak. (B) Effect of increasing the kinesin-1 detachment force parameter was to moderately diminish the minus-end velocity peak. (C) Effect of increasing the kinesin-1 reattachment rate was to diminish the minus-end velocity peak. (D) Effect of decreasing the kinesin-1 stall force was to shift the central velocity peak leftward to better match the experimental data.

The other way to decrease the fraction of the time Kin-DDB complexes are moving solely by DDB is to increase the kinesin-1 reattachment rate. In the ‘basic’ model, we set a constant reattachment rate, *k*_*reattach*_, of 5 s^-1^ for both motors, based on the literature (28,53). To determine whether the kinesin-1 reattachment rates were contributing to the simulation-experiment mismatch, we tested several larger values of *k*_*reattach*_for each motor. In contrast to DDB where altering the reattachment rate had little effect (Fig. 3D), increasing *k*_*reattach*_for kinesin-1 both strongly diminished the fraction of time the complex moves at the unloaded DDB speed and increased the fraction of time that both motors are engaged, resulting in an enhanced cargo velocity peak near zero (Fig. 4C).

The final kinesin parameter we investigated was the kinesin stall force, *F*_*stall*_. In our ‘basic’ model, the kinesin-1 stall force was set to 8 pN and the DDB stall force was set to 3.8 pN based on published optical tweezer experiments (33,46,47). Because of this discrepancy, it is perhaps not surprising that when both motors are engaged, the kinesin directionality dominates. Thus, we investigated kinesin stall forces between 4 and 8 pN, which are in the range of published studies (29,46,60–64), and found that weaker kinesin stall forces cause the central velocity peak to shift closer to zero, better matching the experiments (Fig. 4D).

In summary, a major discrepancy between the experimental and simulated velocity distributions is the peak in the simulations near the unloaded DDB velocity. We found that this minus-end velocity peak could be diminished by decreasing the kinesin-1 unloaded detachment rate or the sensitivity of detachment to load, or by increasing the kinesin-1 reattachment rate. Additionally, decreasing the kinesin-1 stall force resulted in a leftward shift of the dominant peak in the velocity distribution, bringing the simulations in better alignment with experiments.

### Tuning parameters in an improved bidirectional transport model

The simulation results to this point showed that altering motor attachment and detachment rates can decrease the weight of the instantaneous velocity distribution corresponding to the two unloaded motor velocities, and that bringing the kinesin and DDB stall forces into closer alignment can shift the location of the central peak. Informed by these single parameter adjustments, we next formulated an updated model that incorporated multiple parameter adjustments, with the goal of more closely matching the simulated and experimental instantaneous velocity distributions. We chose not to modify the *k*^6^ parameter for either kinesin-1 or DDB because single-molecule TIRF measurements from many labs have consistently measured unloaded detachment rates (equal to velocity divided by run length) near 1 s^-1^ for kinesin-1 and s^-1^ for DDB (29,40,46,47). Our adjustments to kinesin-1 were guided by recent work by Pyrpassopoulos et al. (58), who showed that kinesin-1 KIF5B under purely horizontal loads, achieved through a three-bead optical tweezer geometry, has a ∼6 pN stall force, a 1.1 s engaged duration (matching the unloaded run duration) (58), and a component of its reattachment rate at 100 s^-1^ (31,58). Thus, we set the kinesin-1 stall force to 6 pN and applied an ideal-bond detachment model by setting *k*_*detach*_(*F*) = *k*^6^ . We also increased the kinesin-1 reattachment rate to 50 s^-1^; a 100 rate s^-1^ was not used because there was little improvement beyond 50 s^-1^ and because in the experiments there were components slower than 100 s^-1^ (58). There is no comparable three-bead study for DDB, but we hypothesize that removing the vertical force component may have similar effects on dynein. Thus, we also incorporated a load-independent detachment rate (ideal bond) for DDB.

The parameter adjustments were incorporated into an upgraded model that we named the ‘better-fit’ model (Table S2). Simulations showed that in the ‘better-fit’ model the plus-end side peak was eliminated, the minus-end side peak was diminished, and the central peak was substantially enhanced but was still right-shifted compared to the experimental data (Fig. 5A). Starting from this improved model, we tested further parameter adjustments, with the goal of shifting the central velocity peak to the left and eliminating the minus-end side peak. We first tested the DDB stall force and found that increasing it to 6 pN to match that of kinesin-1 substantially shifted the main peak leftward. Next, we varied the DDB backward stepping rate and found that decreasing it to 3 s^-1^ or 1 s^-1^ also substantially shifted the main peak leftward (Fig. 5C and S3C).

**Figure 5.**
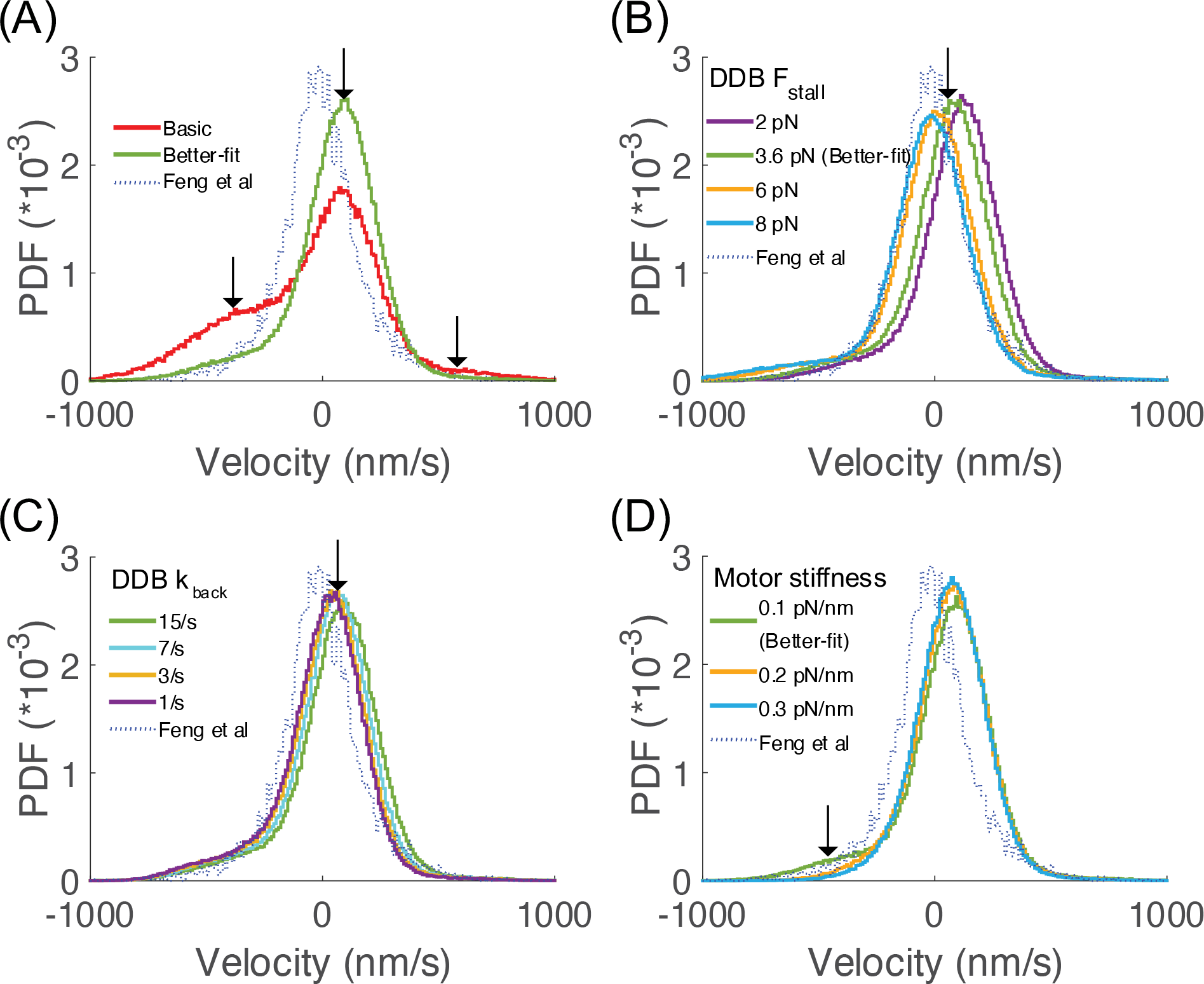
Results from ‘better-fit’ model. (A) Comparison between ‘basic’ and ‘better-fit’ models. The ‘better-fit’ model includes load-independent detachment kinetic for both motors, a 6 pN stall force for both motors, and a 50 s^-1^ reattachment rate for kinesin. Arrows denote the three velocity peaks that are altered. (B) Effect of altering the DDB stall force showed the leftward shift for larger stall forces. (C) Effect of decreasing the DDB backstepping rate is to shift the central velocity peak leftward. (D) Effect of increasing the motor stiffness was to eliminate the small minus-end velocity side peak that results from a motor ‘recoil’ effect at lower stiffness values.

The final property we tested was the motor stiffness, which affects how rapidly motors ramp up their force as they step in opposite directions. We found that increasing motor stiffness eliminated the fastest minus-end velocities in the distribution, and that this weight was shifted to the central peak, bringing it closer to the height of the experimental peak (Fig. 5D and S3D). Upon closer examination, we surmised that this shift results from eliminating a ‘recoil effect’ occurring at lower stiffness values, in which the kinesin detaches and the cargo rapidly moves toward the minus-end to become centered on the engaged DDB motor. A 6 pN force is expected to stretch the spring that connects each motor to its cargo by 60 nm. A recoil of this magnitude over the 0.1 s window used for the velocity distributions corresponds to a -600 nm/s velocity, and this is the component that is eliminated when motor stiffness is increased (Fig. 5D). In summary, in the ‘better-fit’ model either increasing the DDB stall force or decreasing the DDB backstepping rate shifted the central velocity peak leftward to better align with the experimental distribution, and increasing the motor stiffness eliminated the remaining minus-end velocity side peak.

### Using machine learning for parameter sensitivity exploration and optimization

Having narrowed the parameter choices by iterative tuning, we next applied an automated parameter optimization method to identify parameter sets that best match the simulations to the experimental velocity distributions. To do this we used the Matlab Bayesian Optimization function ‘bayesopt’, which identifies an optimal parameter set based on an objective function. We defined the objective function as the residual sum of squares (RSS) between the experimental and simulation velocity distributions, and we defined upper and lower search bounds for each parameter (Table 3). In the first round, we allowed seven parameters to be optimized: the detachment forces and reattachment rates for both motors, along with the motor stiffness, DDB stall force and DDB backstepping rate. Because of the interdependency of the parameters, we found that repeated optimization runs achieved similar RSS values using different parameter sets. Hence, in Table 3 we report the range of optimal parameter values for 10 independent optimization runs, with large ranges connoting parameters that either co-vary with other parameters or do not strongly affect the final velocity distribution, and narrow ranges connoting parameters that most strongly determine model performance. As a final step, we fixed the detachment forces and reattachment rates and performed a second round of optimization on the remaining DDB stall force, DDB backstepping rate, and motor stiffness parameters. The results of this final optimization were similar to the first, but the optimal ranges were narrowed (Table 3).

**Table 3.** *Motor parameters estimated by Matlab ‘bayesopt’ parameter optimization function using the limits shown for each parameter. Parameter ranges from 10 independent optimization runs are shown. From the seven parameter fit, four parameters were fixed at the values shown, and the three remaining parameters were then optimized. Final values refer to parameter values chosen for the ‘best-fit’ model*.

| Parameter | Limits | Optimized parameter values |  | Final value |
| --- | --- | --- | --- | --- |
|  |  | 7 parameter fit | 3 parameter fit |  |
| **Kinesin-1 detachment force** | 0.01 - 50 pN | 17 - 49 pN | Ideal bond | Ideal bond |
| **DDB detachment force** | 0.01 - 50 pN | 5 - 49 pN | Ideal bond | Ideal bond |
| **Kinesin-1 reattachment rate** | 0.01 - 100 s<sup>-1</sup> | 30 - 96 s<sup>-1</sup> | 50 s<sup>-1</sup> | 50 s<sup>-1</sup> |
| **DDB reattachment rate** | 0.01 - 100 s<sup>-1</sup> | 0.3 - 86 s<sup>-1</sup> | 5 s<sup>-1</sup> | 5 s<sup>-1</sup> |
| **DDB stall force** | 0.01 - 6 pN | 5.7 - 6 pN | 5.9 - 6 pN | 6 pN |
| **DDB backstepping rate** | 1 - 20 s<sup>-1</sup> | 1 - 6 s<sup>-1</sup> | 1 - 4 s<sup>-1</sup> | 1 s<sup>-1</sup> |
| **Motor stiffness** | 0.01 - 0.3 pN/nm | 0.2 - 0.3 pN/nm | 0.18 - 0.22 pN/nm | 0.2 pN/nm |

The parameter optimization results can be summarized as follows. First, the optimal detachment forces for both motors were relatively high, consistent with the ideal-bond model we implemented in our ‘better-fit’ model. Therefore, in our final ‘best-fit’ model, we set detachment to be independent of load for both motors. Second, consistent with the ‘better-fit’ model, the kinesin reattachment rate converged in a range around 50 s^-1^ for kinesin and around 5 s^-1^ or slower for DDB. Thus, we chose those values for the reattachment rates. Third, the DDB stall force converged near the 6 pN stall force of kinesin-1, supporting the observation in Fig. 5B that matching the stall forces helps to align the central velocity peak with the experiments. Fourth, the DDB backstepping rate converged in a range of 1-5 s^-1^. This rate is slower than the experimental 15 s^-1^ (33), and reiterates the finding in Fig. 5C that slowing DDB backstepping shifts the central velocity peak leftward. Based on the optimal fit with the reduced parameter set, we set the DDB backstepping rate to 1 s^-1^ (see Discussion). Finally, the motor stiffness converged at 0.2 pN/nm, consistent with our finding that a minimum stiffnesses in this range is needed to avoid the ‘recoil’ phenomenon that contributes unwanted minus-end velocities. The final parameter choices that were used for the ‘best-fit’ model are shown in Table 3, and the entire ‘best-fit’ parameter set is provided in Table 1.

Having identified a final parameter set, we characterized the resulting traces and quantified the fit to the experimental data. The displacement versus time traces for the ‘best-fit’ model were smoother by eye than the ‘basic’ model (Fig. 6A; compare to Fig. 2B, and Fig. S4A; compare to Fig. 1D), and similar to the experimental data (Fig. 2A). Using a mean squared displacement analysis, the apparent diffusion coefficient dropped from 11,800 *nm*^2^*s*^,1^ in the ‘basic’ model to 1272 nm^2^s^-1^ in ‘best-fit’ model, which was close to the 996 *nm*^2^*s*^,1^ for the experimental data (Fig. S1B). More importantly, in the instantaneous velocity distribution (Fig. 6B), the unloaded velocity peaks, which were a significant discrepancy in the ‘basic’ model, were nearly eliminated, and the central peak was shifted to near zero to match the experimental data. One metric of this agreement is that in the ‘best-fit’ model, 95% of instantaneous velocity values fall between -349 to +299 nm/s, similar to the -394 to +371 nm/s range for the experimental data. Fitting the ‘best-fit’ velocity distribution to a Gaussian mixture model identified a central peak centered at -8 nm/s and comprising 93% of weight, along with two side peaks centered at 452 nm/s and -318 nm/s, comprising 0.5% and 6.5% of the weight, respectively (Table 2). Finally, the residual sum of squares (RSS) between experimental and simulation data dropped from 3.93*10^-5^ in the ‘basic’ model to 2.76*10^-6^ in ‘best-fit’ model (Table 2). Other features were also improved; for instance, the simulated traces for the ‘best fit’ model also had longer durations, more closely matching experiments (Fig. S4). Also, for the ‘best-fit’ model the distribution of velocities computed over each trace matched the experimental values (Fig. 6C).

**Figure 6.**
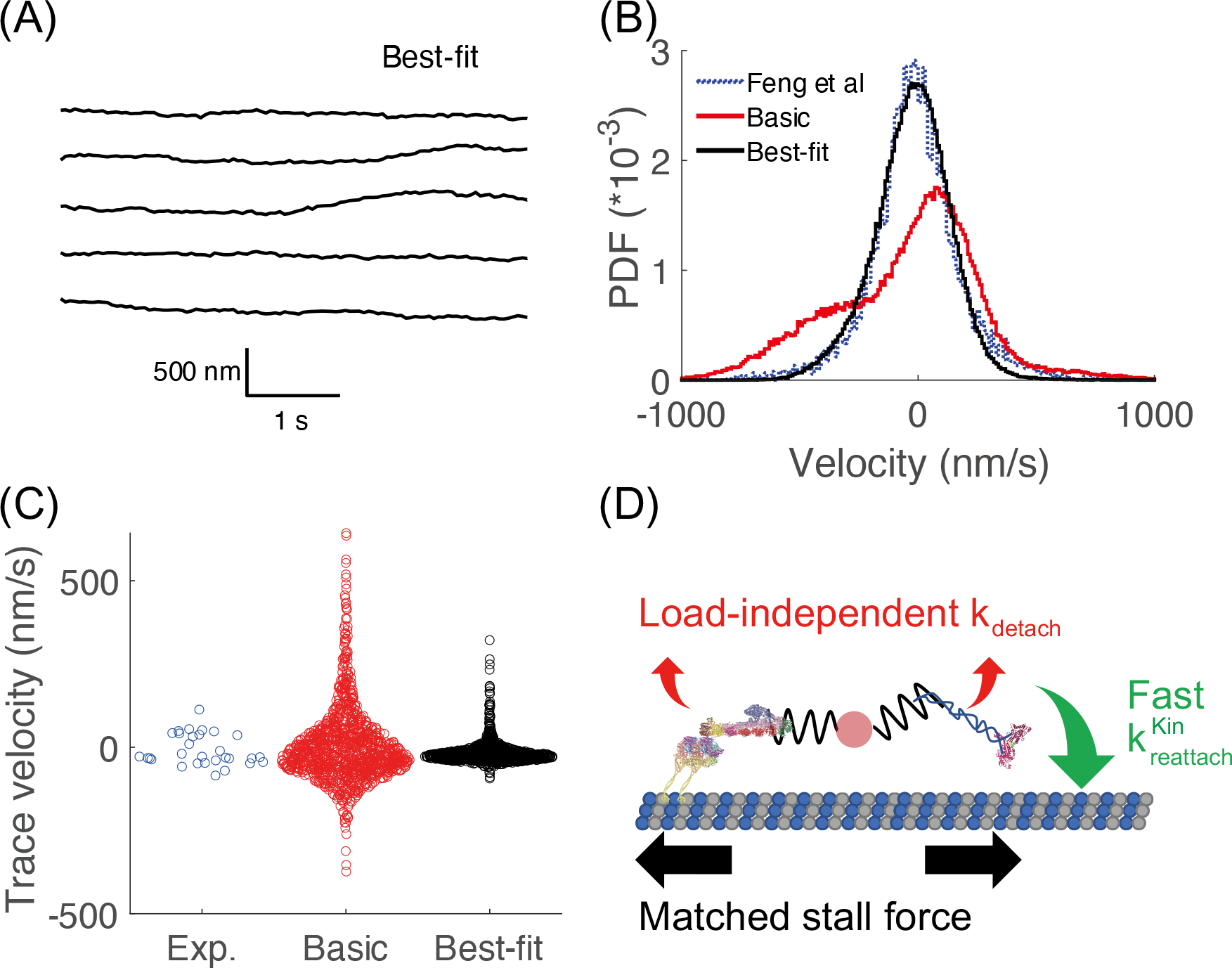
Results from ‘best-fit’ model. (A) Example traces from best-fit model. (B) Comparison of instantaneous velocity distributions between experimental data (blue), basic model (red) and best-fit model (black). (C) Comparison of trace velocity distributions for experimental data (blue), basic model (red), and best-fit model (black). (D) Diagram of the best-fit model for Kin-DDB bidirectional transport, highlighting parameter choices that resulted in optimal alignment with experimental data. The detachment rates of DDB and kinesin are independent of load (ideal-bond), and kinesin-1 reattaches to microtubule with a fast rate (50 *s*^,1^). These features maximize the fraction of time both motors are engaged and pulling against one another. Matching the kinesin and DDB stall forces (not shown) shifted the central velocity peak close to zero to match the experimental data.

## Discussion

Transport of vesicles in axons and dendrites involves varying numbers and types of motors, as well as regulation at multiple levels. Hence, to understand how kinesin and dynein motors attached to the same cargo compete and coordinate during transport, the field has turned to reconstituting the *in vitro* motility of pairs of kinesin and activated dynein (1,7,38,53,65). This approach allows an examination of how differing mechanochemical properties of specific motor isotypes translate into effective movement against an antagonistic partner. Consistent behavior of kinesin-dynein pairs that are observed across different labs (20,32–34) include the following: 1) Kin-DDB complexes remain bound to microtubules for ∼tens of seconds (33,34), considerably longer than kinesin-1 or DDB alone; 2) trajectories include episodes, lasting from seconds to tens of seconds, where the velocity of the complex is ∼10-fold slower than the unloaded velocities of the respective motors; and 3) cargo trajectories are quite smooth and show few if any directional switches. In principle, it should be possible to recapitulate these motility behaviors using Kin-DDB simulations that incorporate established parameters for kinesin and dynein behavior in isolation. However, we find this not to be the case. Instead, simulations of Kin-DDB transport in our ‘basic’ model show frequent directional switches and significant proportions of time when the complexes move at the unloaded velocity of the motors (Fig. 2B). The discrepancies between the simulations and experiments were clearly seen by comparing the instantaneous velocity distributions (Fig. 2C). The experimental data were centered in a wide peak around zero velocity, whereas the simulated data had additional peaks corresponding to the unloaded kinesin-1 and DDB velocities, along with a central peak centered around a slow plus-end speed. Thus, we focused our in silico ‘experiments’ on identifying parameter adjustments that reconciled the simulations with the experiments.

The first modification that helped to align the simulation results with the experiments was to make the kinesin detachment rate independent of load in the range of the stall forces used. Although single-bead optical tweezer experiments have found that kinesin detachment rates increase exponentially with force (58,59,66), recent theoretical and experimental work has suggested that forces perpendicular to the microtubule inherent to the geometry of these experiments may be contributing to the detachment rate. In support of this, experiments using a three-bead assay that almost fully eliminate these vertical forces found that the kinesin-1 detachment rate was approximately independent of load (58). Incorporating this ‘ideal-bond’ for kinesin into our model strongly diminished the minus-end side peak in the velocity distribution; thus, our simulations support the hypothesis that against forces oriented purely parallel to the microtubule, the kinesin detachment is independent of load. We also examined a catch-bond model for kinesin, and found a further increase in the amplitude of the central velocity peak with increasing catch bond properties (Fig. S5). Thus, our results are consistent with possibility that kinesin has catch-bond behavior under some circumstances, but an idea-bond for kinesin was sufficient in our simulations to match the experimental data.

We also found that incorporating a load-independent detachment rate for DDB diminished the fast plus-end velocities seen in the simulations, better matching the experimental data. This result can be understood simply as DDB remaining bound to the microtubule against the pulling forces of kinesin, thus preventing durations in which DDB is detached and kinesin is moving at its unloaded speed. A DDB ideal-bond model is supported by experiments with activator-free dynein and with isolated vesicles, which found evidence that dynein has catch-bond behavior under some conditions (29,57), though we note that a DDB optical trapping study by Elshenawy et al. found an exponential dependence of DDB detachment with load (33). However, it remains a possibility that, as has been shown for kinesin-1 (30,33,59), vertical forces inherent to the single-bead optical trapping assay may contribute to detachment of DDB. Thus, our simulations put forth the testable hypothesis that when pulling exclusively against hindering loads oriented parallel to the microtubule, DDB detachment is independent of load over the forces generated by a single kinesin motor. As with kinesin, our results were also consistent with DDB catch bond behavior (Fig. S3A), but an ideal bond was sufficient to match the simulations to the experiments.

Incorporating a fast kinesin-1 reattachment rate was another important modification that helped to bring the simulations into alignment with the experimental data. In particular, the fast kinesin-1 reattachment rate diminished the minus-end velocity side peak, which can be understood as minimizing the time that DDB moves at its unloaded velocity. Our basic model used a kreattach value of 5 s^-1^, which was initially determined in a study that measured motor-driven deformations of giant unilamellar vesicles (53), was later supported by experiments that used DNA to connect two kinesins (48), and which has been employed in a number of modeling studies (e.g. (28)). However, three recent optical tweezer studies found that against hindering loads, kinesin-1 can slip backward in multiple 8 nm intervals and rapidly reengage with the microtubule at rates consistent with our 50 s^-1^ reattachment rate (58,67,68). These experiments suggest that against a hindering loads oriented parallel to the microtubule, kinesin-1 enters a weak-binding state in which it slips backward, and then rapidly reestablishes a strong-binding state. We also note that this 50 s^-1^ reattachment rate is still below the 125 s^-1^ reattachment rate predicted from multiplying the bimolecular on-rate of kinesin in solution by the effective tubulin concentration when two motors are connected through a DNA linker (48). Thus, our simulations put forth the testable hypothesis that when pulling against hindering loads oriented parallel to the microtubule, the kinesin-1 reattachment rate is 50 s^-1^ or faster. A possible mechanism to explain this phenomenon is that if a kinesin slips or detaches against a load oriented parallel to the microtubule, the motor is presented with multiple sites of reattachment as it slides backward. Further, it is possible that if the motor reattaches against a hindering load, the force enhances the rate of ADP release, which maximizes the probability that the motor enters a strongly-bound state and restarts its walking cycle.

The model adjustment that most influenced the position of the central peak in the instantaneous velocity distribution was altering the relative stall forces of DDB and kinesin-1, highlighting that the antagonistic motors rapidly establish a “draw” where each is pulling at near its maximum force. The paired modification of reducing the kinesin-1 stall force from 8 pN down to 6 pN and increasing the DDB stall force from 3.6 pN up to 6 pN shifted the central peak leftward by roughly ∼140 nm/s to match the experimental peak (Fig. 6B). A large body of experiments support a kinesin-1 stall force in the range of 6 pN (29,46,60–64), justifying this modification. For DDB, one justification for increasing the stall force is that the lower value we started with was taken from a study using single-bead tweezer geometry (33), and it is possible that in the geometry of these DDB-kinesin pairs where forces are oriented solely parallel to the microtubule, the DDB stall forces are closer to the ∼6 pN of kinesin-1 (Fig. 3D). Related to the stall force, we also found that slowing the DDB backstepping rate shifted the central velocity peak leftward. This shift makes sense in cases where the kinesin stall force is greater than the DDB stall force, meaning that DDB is under superstall forces and stepping backward at 15 s^-1^ (Fig. 5C and S3C). However, we note that when stall forces were matched, the DDB backstepping rate did not have a strong effect on the velocity distribution (Fig. S6). Hence, although single-bead optical tweezer experiments measured a DDB backstepping rate of 15 s^-1^ (33), our simulations put forward the testable hypothesis that the DDB backstepping rate may be smaller in the absence of any vertical forces.

Another model adjustment that had a surprising effect was increasing the stiffness of the linkages connecting the two motors to the cargo. When model and experiments were compared by analyzing instantaneous velocities, a ‘recoil’ effect becomes apparent following detachment of one of the motors (usually kinesin in our simulations). Thus, the simulations provide a lower limit of 0.2 pN/nm for the motor stiffness and make the testable prediction that if a more compliant DNA linkage were used to connect the motors, the instantaneous velocity distribution would reveal apparent fast velocities due to large displacement recoil events. Additionally, a more compliant linkage between the motors would be relevant to the transport of large (0.1 – 1 micron) vesicles in cells, where motor forces are sufficient to deform the membrane.

Despite the large body of single-molecule data describing their motor properties, it remains challenging to predict the bidirectional dynamics that will result from antagonistic kinesin and dynein motor pairs. In the present work we find that by using experimental data to constrain model simulations, mechanistic insights can be generated into how kinesin and DDB operate in antagonistic pairs. Our simulations support a model (Fig. 6D) in which: 1) Over the range of forces generated by the motors, the detachment rate of both kinesin-1 and DDB is insensitive to load; 2) In the two-motor geometry examined, the kinesin-1 and DDB stall forces are similar; 3) kinesin-1 reattaches to the microtubule at 50 s^-1^ in the geometry of these motor pairs; and 4) when modeled as linear springs, motor stiffness is at least 0.2 pN/nm. These model- generated hypotheses generate testable predictions for future single-molecule experiments. Furthermore, by identifying motor parameters that are the strongest determinants of bidirectional transport, this work provides a framework to interpret how diverse kinesins, different activating dynein adapters, and motor- cargo stiffness will affect the resulting bidirectional transport of intracellular cargo.

## Author Contributions

T-C.M. and W.O.H. designed research, T-C.M. performed all simulations and carried out analysis, Q.F. and A.M.G. carried out experiments and experimental data analysis. T-C.M., W.O.H. and A.M.G. wrote the manuscript, and all authors edited the manuscript.

## Acknowledgements

The authors thank George Ohashi and John Fricks who helped formulate a previous iteration of the current model, as well as members of the Hancock lab who provided helpful input. We thank an anonymous reviewer for suggesting an objective parameter optimization strategy. This work was supported by NIH grants R01GM076476, 1R01GM122082 and R35GM139568 to W.O.H., and F32GM137487 to A.M.G.

## Supplemental Figures and Tables

**Figure S1.**
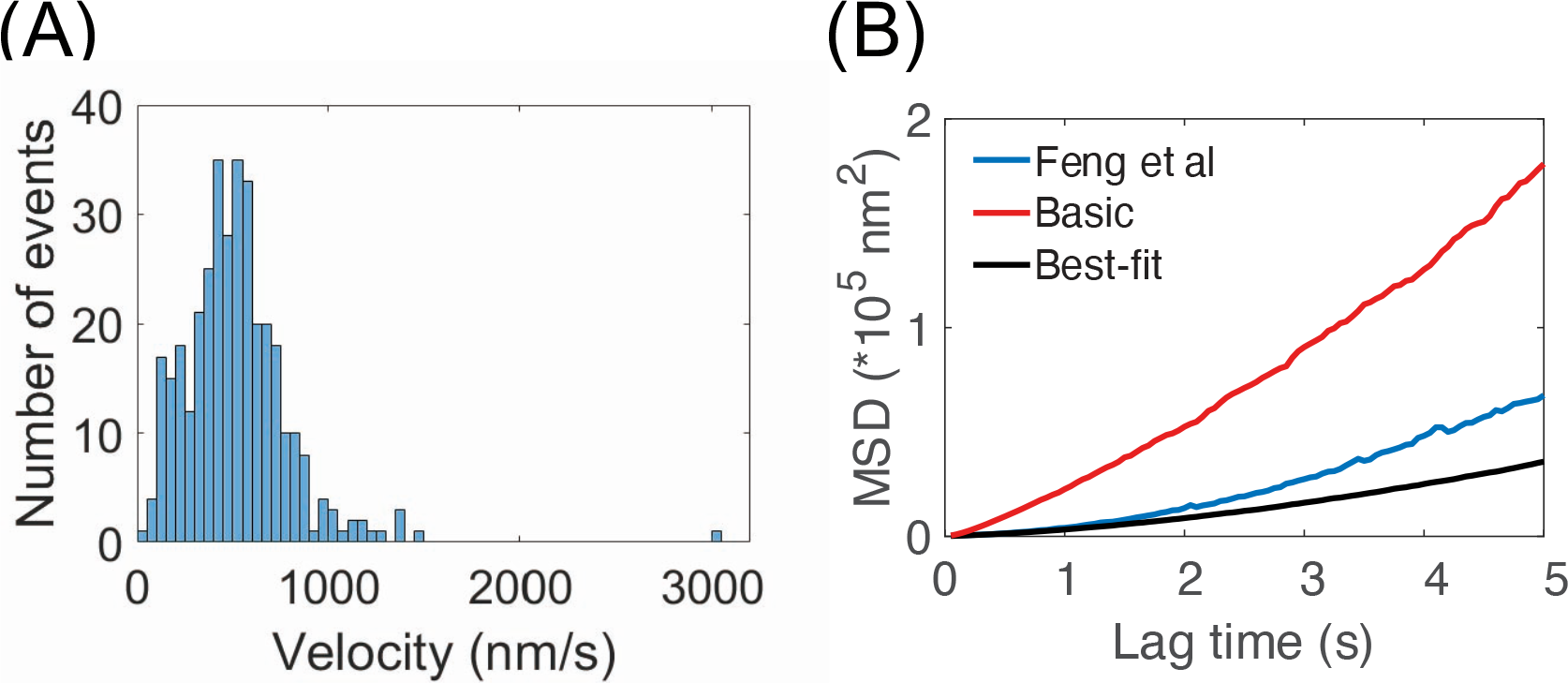
(A) Distribution of trace velocities of isolated kinesin-1 in the dynein motility buffer used in Kin- DDB experiments (1). Dynein motility buffer consists of 30 mM HEPES, 50 mM potassium acetate, 2 mM magnesium acetate, 1 mM EGTA and 10% glycerol, supplemented with 2 mg/ml casein, 20 mM glucose, 37 mM βME, glucose oxidase, catalase, 10 mM Taxol, and 2 mM ATP. The mean velocity of 515 nm/s was used for the unloaded kinesin-1 velocity in the simulations. (B) Mean squared displacement (MSD) analysis of experimental data from Feng et al. (1) and Basic and Best-fit model results from this study. Curves were fit with the equation: *MSD* = *V*^2^*t*^2^ + 2*Dt*, where V is velocity, D is the apparent diffusion coefficient, and t is the lag time. Fit results are *V* = 49 *nm*/*s*, *D* = 996 *nm*^2^/*s* for experimental data, *V* = 49*nm*/*s*, *D* = 11,800 *nm*^2^/*s* from the basic model, and *V* = 32 *nm*/*s*, *D* = 1812 *nm*^2^/*s* from the best-fit model.

**Figure S2.**
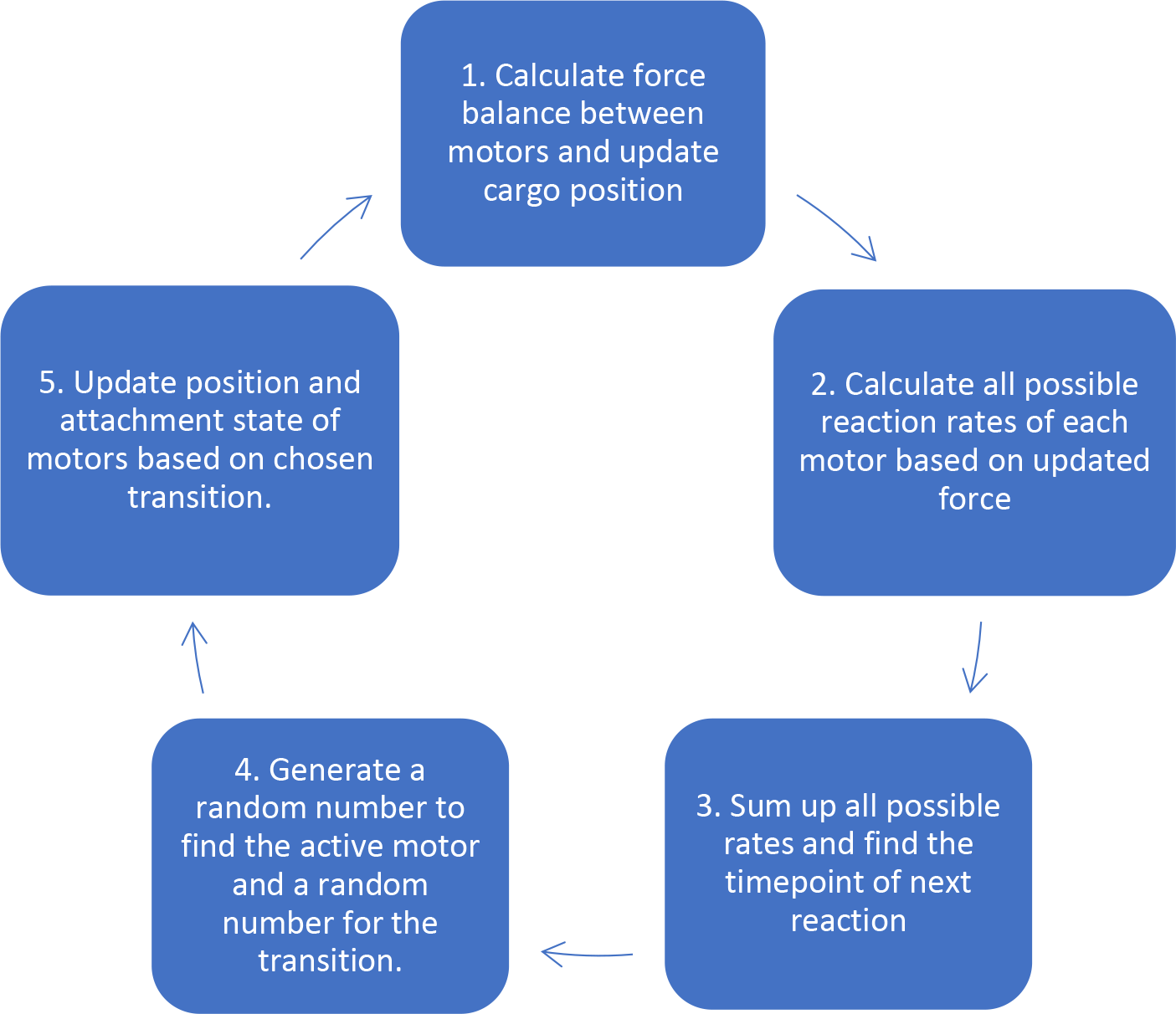
Flow chart for stochastic bidirectional stepping algorithm. The initial positions of both motors and the cargo are at the origin. The first step is to update the force balance between the motors and use this do define the cargo position. The second step is to calculate all possible reaction rates based on the forces applied to each motor. The third step is to use a random number to find the timepoint when the next reaction occurs based on Eq. (2) in Methods. The fourth step is to use one random number to choose which motor is active and a second random number to choose which transition the active motor will make. Finally, the motor position and attachment states are updated. The steps are repeated until both motors detach from the microtubule or until the maximum simulation time is achieved.

**Figure S3.**
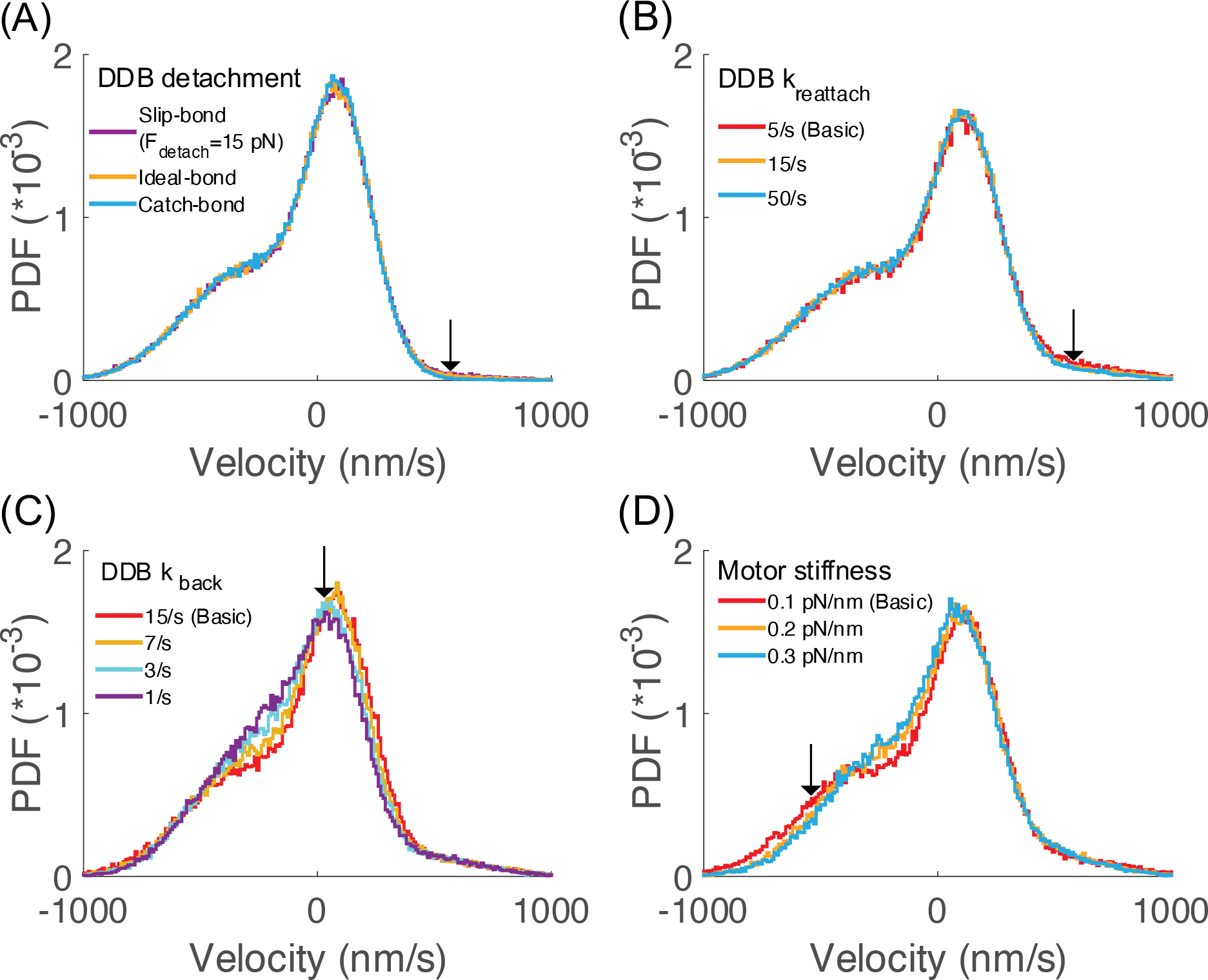
Effect of changing DDB mechanochemical properties and motor stiffnesses on the instantaneous velocity distribution in the ‘basic’ model. (A) Comparison between different load- dependencies of the DDB detachment rate. The slip-bond with Fdetach = 15 pN was taken from Fig. 3C, an ideal-bond corresponds to Fdetach set to infinity, and the catch-bond used Fdetach = -3 pN such that detachment slows with increasing force. (B) Effect of altering the DDB reattachment rate, showing only a very small effect on the plus-end side peak. (C) Effect of altering the DDB backstepping rate, *k*_*back*_, showing a small leftward shift of the central velocity peak with slower backstepping. (D) Effect of increasing the motor stiffness was to reduce the minus-end velocity peak.

**Figure S4.**
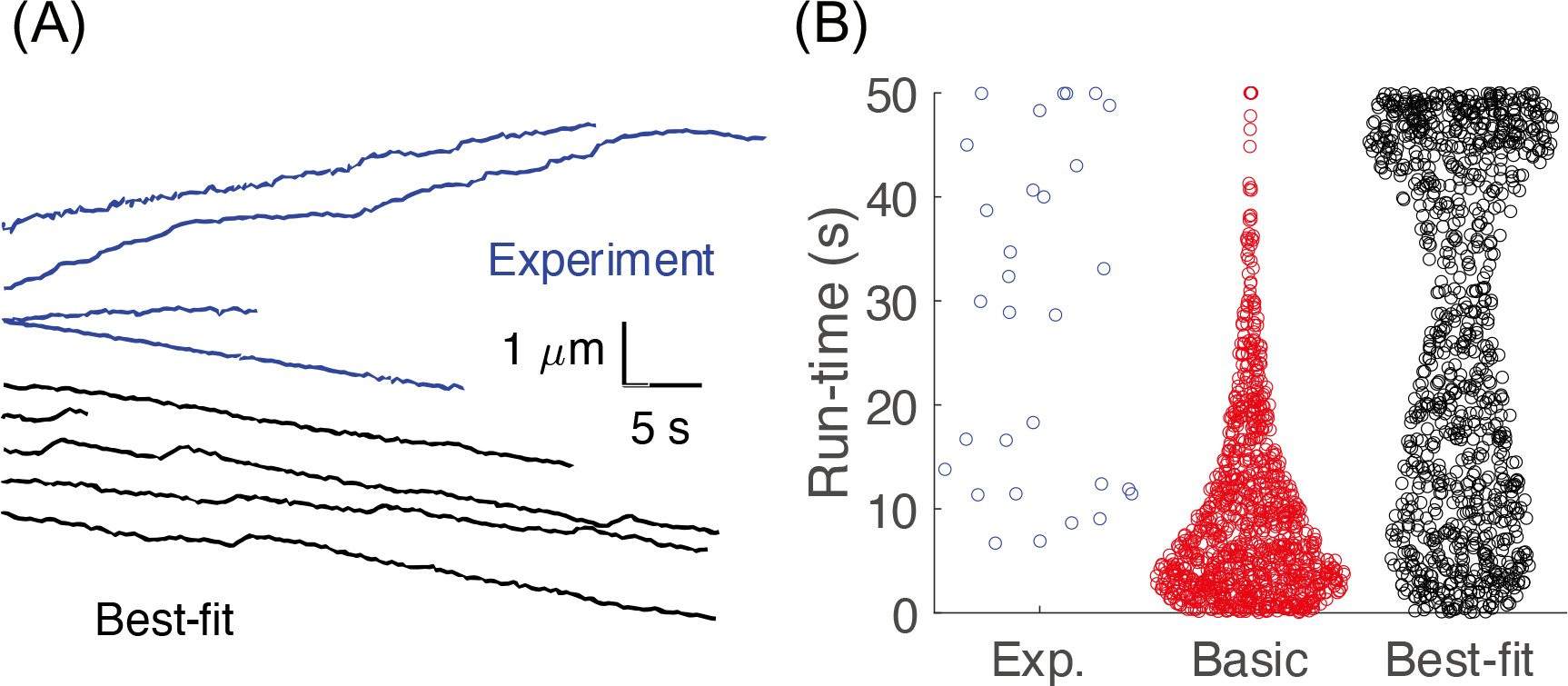
(A) Example 50-second traces from tracking experiment (1) and simulations of the best-fit model. (B) Distribution of run durations from experimental data (blue), basic model (red) and best-fit model (black). The best-fit model had longer run durations that more closely matched the experimental data. Maximum duration of 50 s was set by movie duration from experiments.

**Figure S5.**
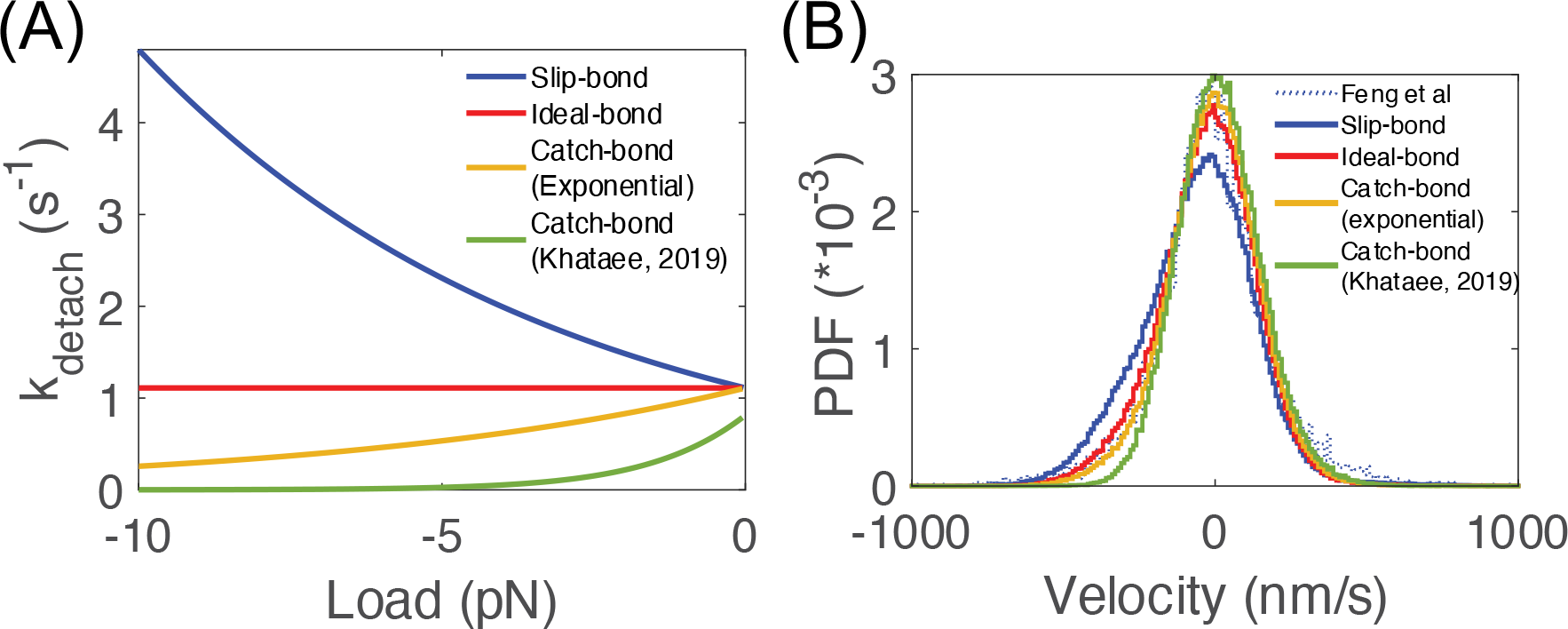
(A) Different models of kinesin detachment under horizontal hindering loads. The slip-bond model (blue) is based on experimental data from Andreasson et al. (2). The ideal-bond model (red) used a load-independent detachment rate, *k*_*detach*_(*F*) = *k*^0^_*detach*_, where *k*^0^_*detach*_ is unloaded detachment rate. The exponential catch-bond model (yellow curve) reversed the sign of the *F*_*detach*_ = 6.8 pN used for the slip bond, such that 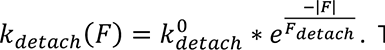. The alternate catch-bond model (green) is the detachment rate under horizontal load calculated by Khataee et al. (3). (B) The instantaneous velocity distribution of ‘best-fit’ kin-DDB model with different kinesin detachment models applied.

**Figure S6.**
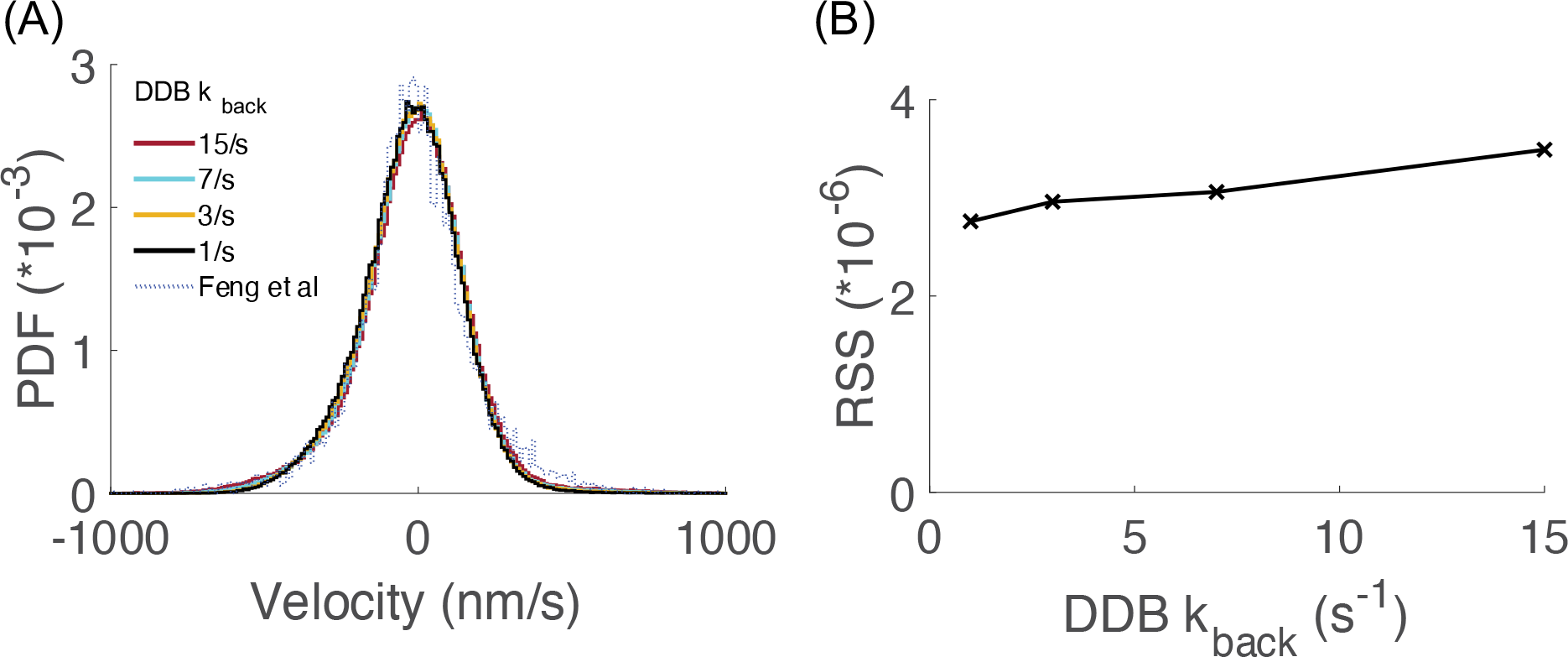
Effect of changing the DDB backstepping rate in the ‘best-fit’ model. (A) Instantaneous velocity distribution, showing the small effect of changing the DDB backstepping rate. (B) Residual sum of squares between simulation and experiment for ‘best-fit’ model with different DDB backstepping rates.

**Table S1.**
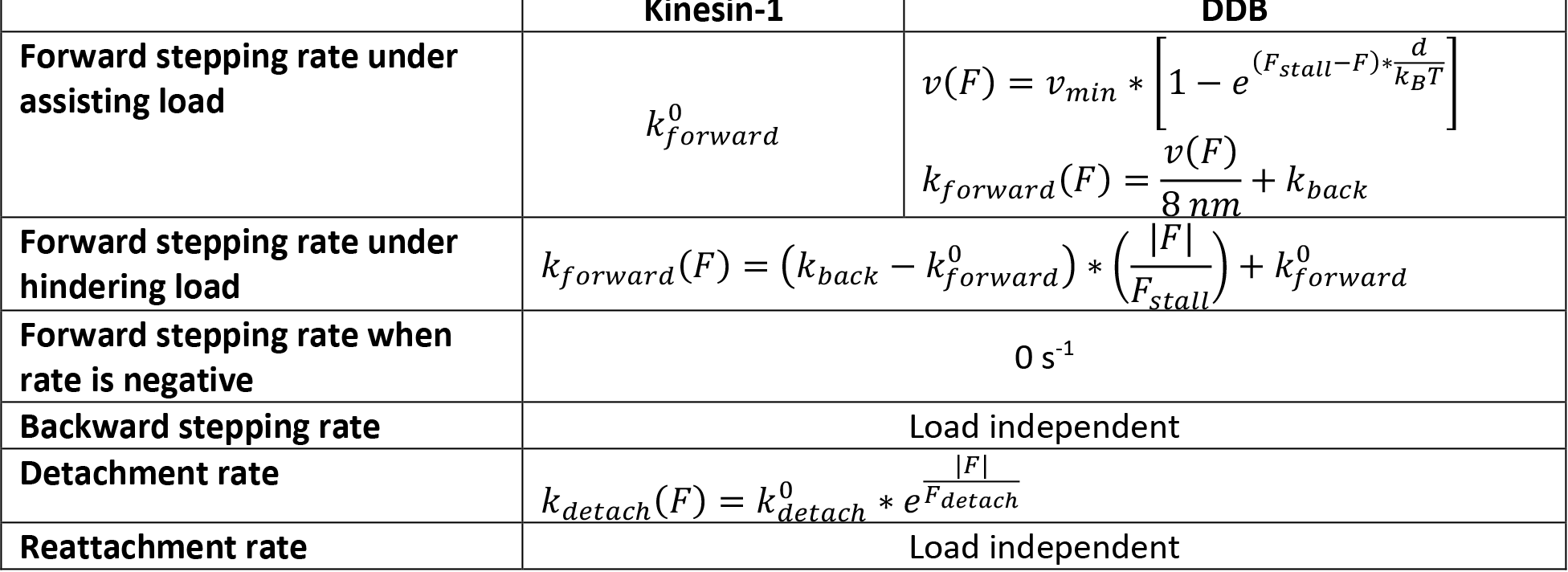
*The models for load-dependent motor mechanisms used in stochastic stepping model*.

**Table S2.** Parameters used for the ‘better-fit’ model. ^a^ For kinesin under assisting loads, the unloaded detachment rate extrapolation was 7.4 s^-1^ and detachment force was 12.8 pN based on (2). For DDB under assisting loads, the unloaded detachment rate and detachment force were identical to the hindering load condition. Ideal bond (load-independent detachment) corresponds to Fdetach equal to infinity. ^b^ The parameters used in “better-fit” were established by results of parameter sensitivity tests in simulation.

| Parameter | Better-fit model |  |
| --- | --- | --- |
|  | Kinesin-1 | DDB |
| Unloaded velocity (nm/s) | 515 | 328 |
| Backstepping rate ($s^{-1}$) | 3 | 15 |
| Stall force (pN) | 6<sup>b</sup> | 3.6 |
| Unloaded detachment rate ($s^{-1}$)<sup>a</sup> | 1.11 | 0.1 |
| Detachment force (pN) | Ideal | Ideal |
| Reattachment rate ($s^{-1}$) | 50<sup>b</sup> | 5 |
| Stiffness (pN/nm) | 0.1068 | 0.1068 |
| Residual sum of squares (RSS) | 4.29*10<sup>-5</sup> |  |

